# Superior colliculus neurons encode and causally shape somatosensory decisions

**DOI:** 10.64898/2026.07.01.735970

**Authors:** Saba Gharaei, Matthew F. Tang, Greg J. Stuart, Ehsan Arabzadeh

**Author notes:** Co-senior authors (Greg J. Stuart, Ehsan Arabzadeh).

## Abstract

The ability of an organism to attend to, and orient towards, relevant stimuli is critical for survival. In the mammalian brain, a principal brain region performing this function is the superior colliculus (SC). Despite its important role in attention and orienting movements, little is known about the role the SC plays in sensory decisions. Using Neuropixel recording and optogenetic perturbation of neuronal activity in awake behaving mice, we provide a quantitative link between the activity of neurons in the SC and the behavioural outcome during a whisker-dependent sensory detection task. Consistent with the idea that the SC contributes to sensory decision-making, the activity of SC neurons was correlated with behavioural performance. Furthermore, we found that optogenetic inhibition of the SC reduced behavioural performance without reducing sensitivity or reaction time. These findings indicate that the SC both encodes and causally shapes sensory decisions during a whisker-dependent task.

## Introduction

The ability of an organism to attend to, and orient towards, threats is critical for survival. For example, when we cross the road, our brain integrates information across multiple sensory modalities, selects which parts of the oncoming traffic we need to attend to and ultimately decides when it is safe to cross. Similarly, prey animals (like rodents) need to detect predators, integrate sensory information from the environment, and take evasive action. A key brain area thought to be involved in this process is the midbrain structure called the superior colliculus (SC). Consistent with this idea, the SC receives direct input from multiple sensory modalities and plays an important role in moving the eyes, head and body towards (or away from) biologically significant stimuli ^1–6^. In addition to direct subcortical input, the SC also receives sensory information from the primary sensory cortices ^7–11^. To what extent sensory information to the SC contributes to sensory-driven decision-making is unclear and is the focus of this study.

Although historically framed as a reflexive orienting centre, the SC can also modulate sensory responses in primary and higher-order cortex ^12–20^, indicating a broader role in attention and decision-related computations. In addition, other research indicates that the SC integrates sensory input with behavioural goals to guide selection of behaviourally relevant stimuli or actions ^21–26^. Recent work further shows that the deeper regions of the SC integrate sensory input with basal ganglia signals to shape orienting behaviour ^27^, positioning the SC as a key structure linking sensory input to behavioural responses. Together, these data support the idea that subcortical circuits play a role in perceptual choices.

The whisker sensory pathway provides a powerful system in rodents to examine the potential role of the SC in sensory decisions. Rodents rely heavily on their whiskers for navigation and object localisation, and a major part of the somatosensory input to the rodent SC arises from the whiskers ^28–32^. Other work indicates that both GABAergic and non-GABAergic neurons in the SC receive cortical and brainstem whisker input ^31^, which presumably plays an important role in supporting coordination of whisker input and whisker-related motor output. Here, we use the whisker system to directly test the idea that the SC contributes not only to orienting movements and defensive behaviours in mice, but also to the evaluation of sensory evidence during sensory decisions. To do this, we first quantify neuronal activity in the SC during a whisker-based detection task, then use optogenetic inhibition of the SC to establish a causal role of activity in the SC in somatosensory decision-making.

## Results

### Contribution of the superior colliculus to sensory-guided behaviour

Understanding how neuronal activity drives sensory decisions is a fundamental objective in neuroscience ^33–37^. To address this issue, we determined how neuronal activity recorded in the SC relates to behavioural output in a binary Go/NoGo whisker-detection task (Figure 1a). Mice (n=4) were trained to lick for a water reward in response to an air-puff to their whiskers. They learned to respond reliably to the whisker stimulus and showed significantly higher lick rates on stimulus trials (43.1 ± 3.7%) compared to catch (no-stimulus) trials (6.5 ± 1.3%; Figure 1b–f), corresponding to a behavioural sensitivity (d′) of 1.37 ± 0.12 and an AUROC of 0.68 ± 0.02, both significantly above chance (p < 0.001). The mean first lick time was 0.66 ± 0.02 s after stimulus onset (25 sessions). To examine the role of neuronal activity in the SC during this sensory-based decision task, we recorded single-unit activity from the anterolateral SC with Neuropixel 1.0 probes. Figure 1g shows the response of an example SC neuron to whisker stimulation. Population responses from an example session (130 neurons) and across all recordings (2,507 neurons) are shown in Figure 1h.

**Figure 1:**
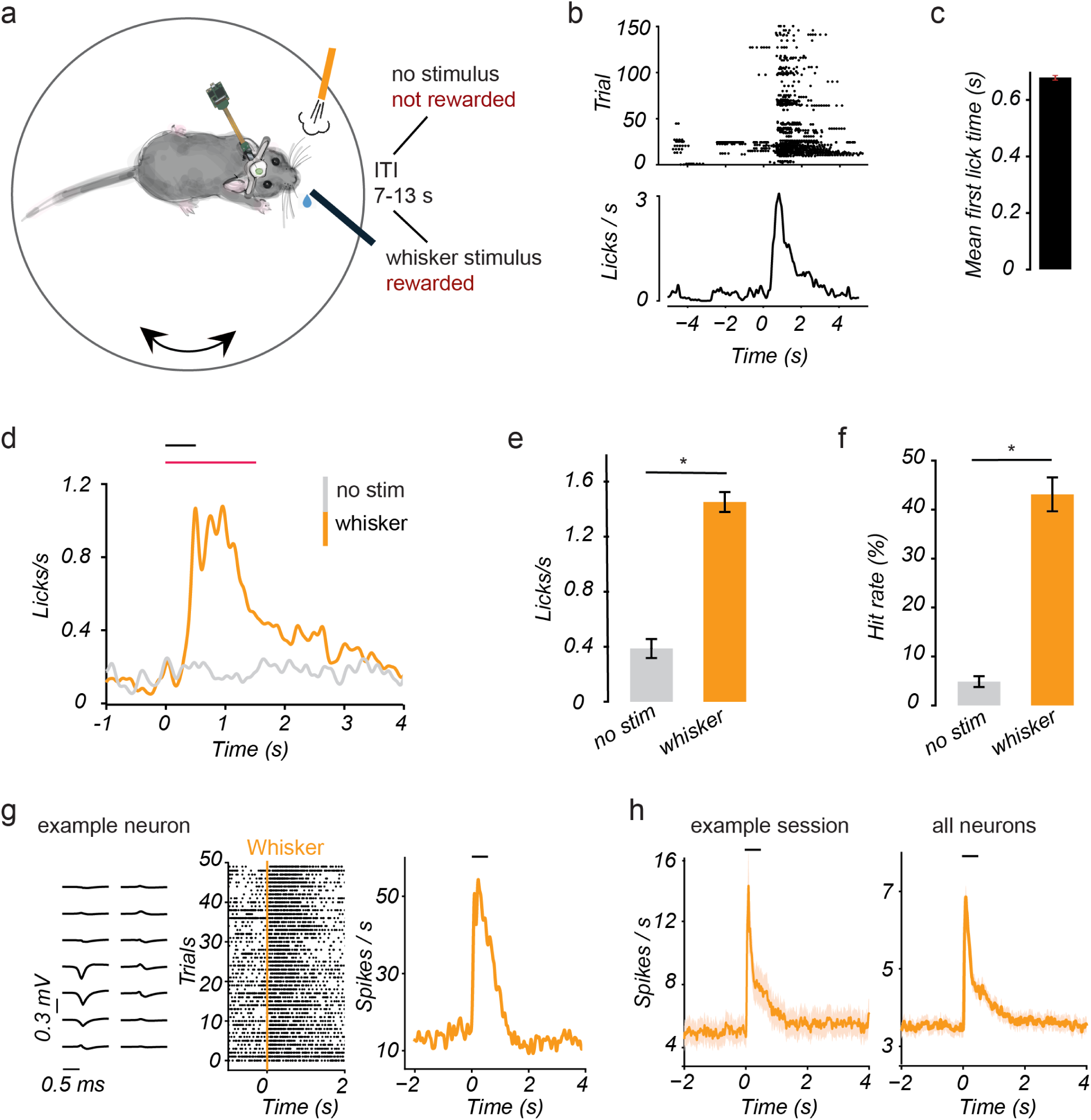
Superior colliculus activity during a whisker-based detection task. **a)** Schematic of the experimental setup. Head-fixed mice were placed on a freely rotating disc while performing a whisker detection task. Licking the spout during the 1.5 s reward window following stimulus presentation resulted in a sucrose-water reward. Each trial was separated by an inter-trial interval of 7–13 s. **b)** Raster plots (top) and peri-stimulus time histograms (bottom) from an example session showing successful task performance; time 0 indicates the onset of the whisker stimulus. **c)** Mean first lick time following whisker stimulation (25 sessions). Error bars represent SEM. **d)** Average lick profile for whisker-stimulus and catch (no-stimulus) trials. Mice licked reliably following whisker stimulation but not during catch trials. The black line indicates stimulus duration (0.5 s), and the red line marks the 1.5 second reward window. **e)** Mean lick rate across sessions (number of animals = 4; total sessions = 25). Asterisk indicates p < 0.05, two-sided paired t-test; Error bars = SEM. **f)** Mean Hit rate across sessions; same conventions as in (e). **g)** Left: spike waveforms of an example SC neuron recorded on overlapping Neuropixel channels. Middle: raster plots showing responses to air-puff whisker stimulation. Right: peri-stimulus time histograms (PSTH) of spiking for the same neuron. **h)** Left: Population PSTH from an example recording session (130 neurons). Right: Population PSTH across all recordings (4 mice, 13 sessions, 2,507 neurons). Shaded areas represent ± SEM.

To dissociate neuronal activity driven by sensory input from movement-related activity, we compared neuronal and lick timings using methods similar to DiCarlo and Maunsell (2005). Across trials, we characterised neuronal responses in relation to either the onset of licking or the onset of the sensory stimulus. In most neurons (77%), the maximum firing rate in a time window (±100 ms) centred around the stimulus onset was higher than the firing rate centred on lick onset (Figure 2a; 1,222 neurons lay below and 359 neurons lay above the unity line, respectively). This indicates that SC activity was more strongly aligned to the stimulus than to lick onset in the majority of neurons. Figure 2b visualises how responses of two example neurons were time-locked to sensory stimulus onset, regardless of when licking occurred. This relationship is further illustrated in the population averages, showing a sharp increase in neuronal responses when activity was aligned to stimulus onset compared to when it was aligned to the time of the first lick (Figure 2c). These observations indicate that neuronal activity was primarily associated with stimulus onset rather than licking. Finally, we examined neuronal responses during spontaneous, non-rewarded licks occurring before stimulus presentation (Figure 2d). This analysis showed that spontaneous licks were not associated with a clear time-locked modulation of neuronal activity, supporting the conclusion that our recordings primarily reflect sensory rather than motor-related responses.

**Figure 2:**
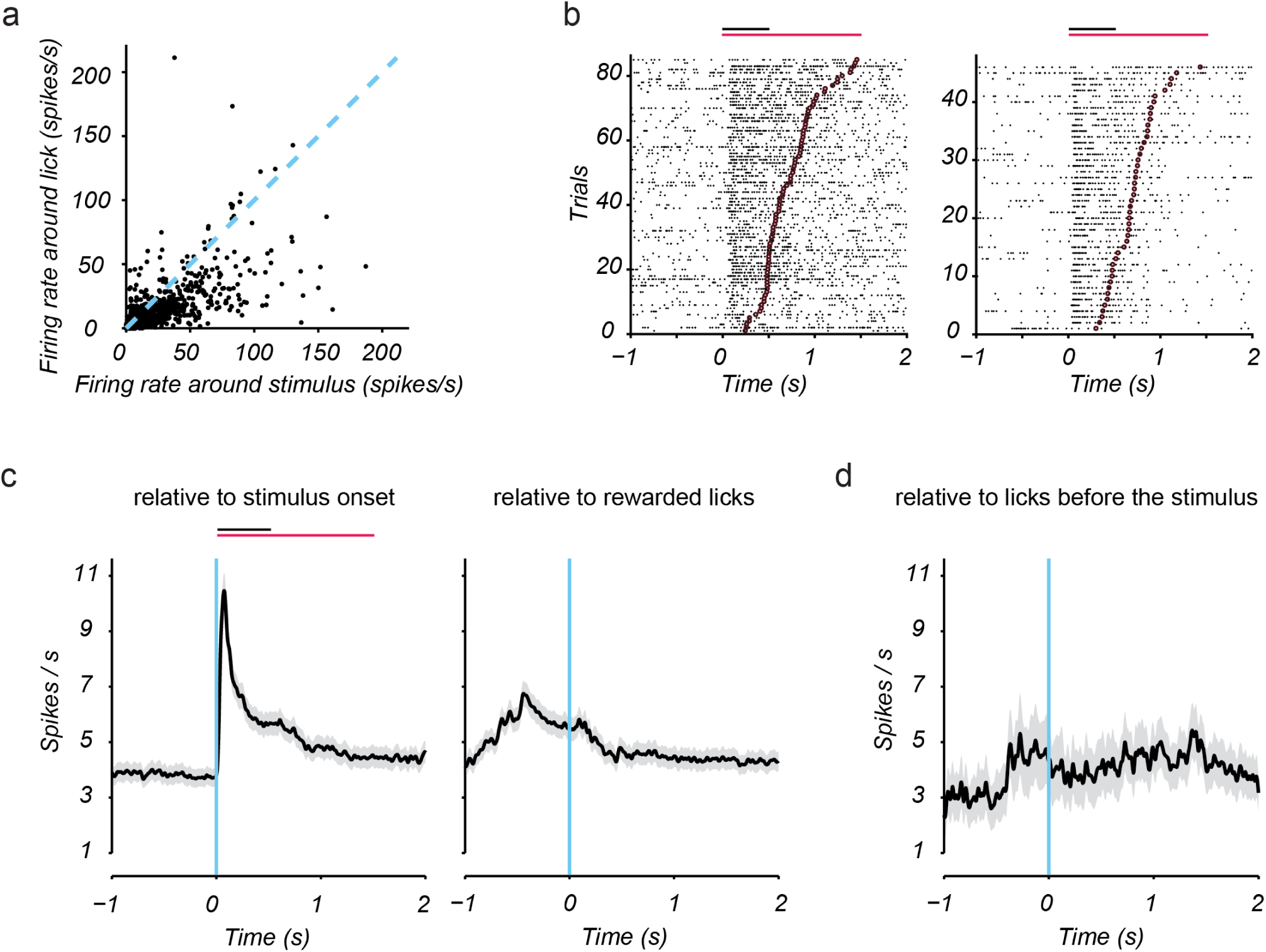
Superior colliculus activity is aligned to sensory input rather than licking behaviour. **a)** For each neuron, we plotted the maximum firing rate in a time window centred on rewarded licks (±100 ms) against the maximum firing rate in a time window centred on stimulus onset (±100 ms). For this comparison, only sessions with at least 35 Hit trials were included (7 sessions; 1,581 neurons). Each dot = one neuron. Most points (n = 1,222) lie below the unity line, indicating stimulus-related responses. **b)** Raster plots for two example SC neurons during Hit trials only. Each black dot = a spike. Time 0 = stimulus onset. Trials are sorted by reaction time (time to first lick; burgundy circles). For both neurons, the response reflects stimulus onset rather than lick time. Black line = stimulus (0.5 s); red line = reward window (1.5 s). **c)** Population spiking activity of SC neurons (n = 1,581) aligned to stimulus onset (left) or first lick (right) across 368 trials. Shaded areas = SEM. **d)** Spiking aligned to spontaneous licks before stimulus onset (50 trials). Shaded areas represent ± SEM.

We next examined whether neuronal activity in the SC correlated with behavioural performance. Based on the analyses described above, we identified 1,222 neurons whose activity was greater just after stimulus onset (within 100 ms) than during licking. We analysed this population to determine how their activity compared during Hit (correct detection) and Miss (failed detection) trials within a 100-ms post-stimulus window (Figure 3). Across the population, the peak sensory-evoked response in these neurons was significantly higher during Hit compared to Miss trials (Figure 3a-c; p < 0.001; two-sided paired t-test). These results indicate that sensory responses in the SC are associated with behavioural outcome. Beyond the increase in response magnitude, the mean spike latency was also significantly shorter during Hit compared to Miss trials (Figure 3d; p < 0.05; two-sided paired t-test), consistent with more rapid and/or more synchronous sensory processing during successful detection and slower, less synchronous processing during unsuccessful detection. We also found that the baseline firing rate (100 ms pre-stimulus) was slightly higher on Hit compared to Miss trials (Figure 3e-f; p < 0.01; two-sided paired t-test), suggesting that the internal state or arousal level may also contribute to behavioural performance.

**Figure 3:**
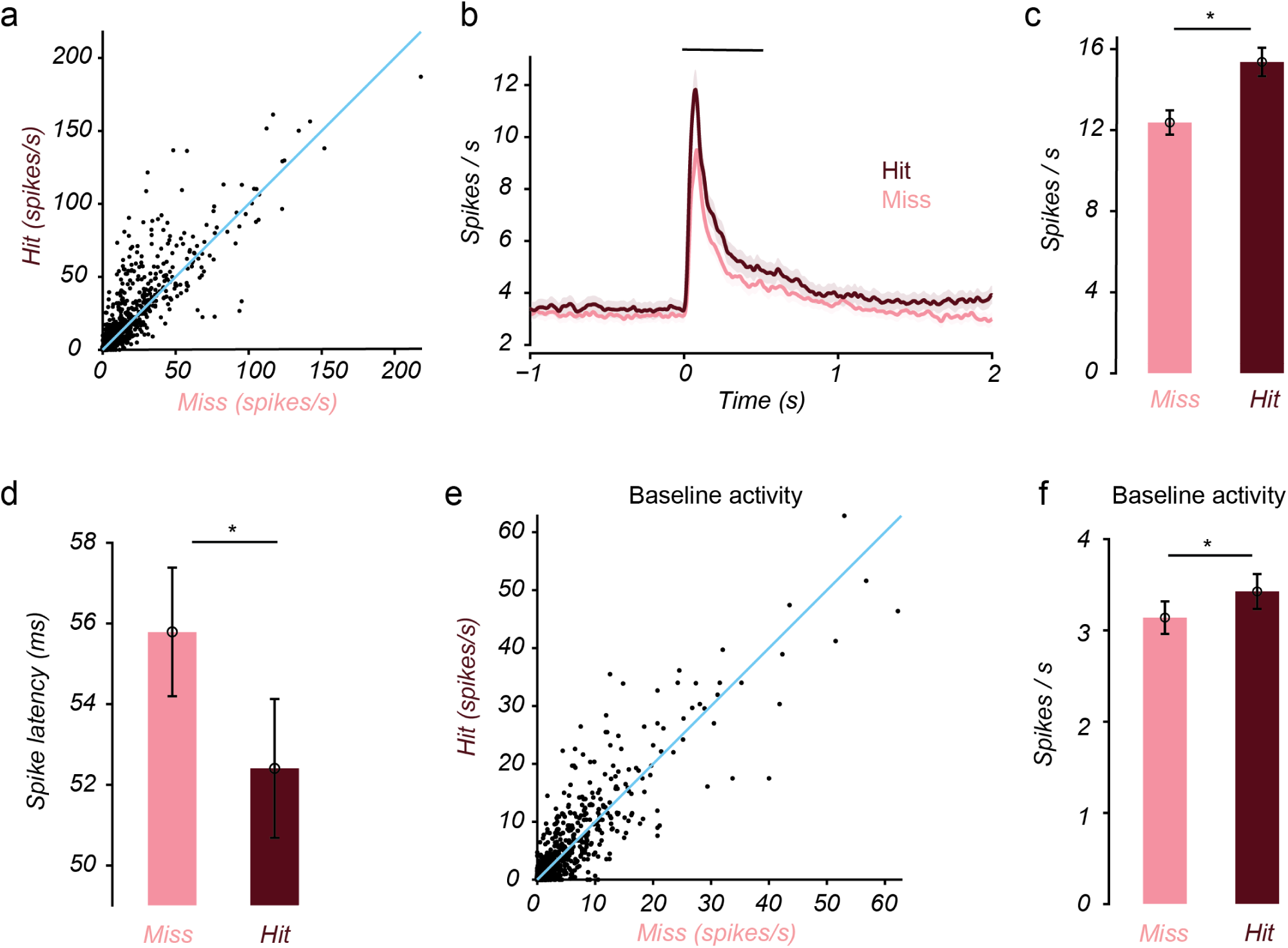
Superior colliculus activity correlates with behavioural outcome. **a)** For each neuron, we plotted the maximum firing rate within the first 100 ms following stimulus onset for Hit and Miss trials. For these analyses, only neurons with greater stimulus-aligned than lick-aligned activity were included (1,222 neurons). **b)** Mean population activity over time for Hit (burgundy) and Miss (pink) trials. Time 0 = stimulus onset; black line = 0.5 s stimulus period. **c)** Bar graph showing the average peak firing rate within the first 100 ms following stimulus onset for Hit and Miss trials. Asterisk indicates p < 0.05, two-sided paired t-test. Error bars = SEM. **d)** Average spike latency for Hit and Miss trials; latency was defined as the time of the first of two consecutive significant 5-ms PSTH bins within the first 200 ms following stimulus onset. **e)** Baseline (100 ms pre-stimulus) activity for Hit vs Miss trials for all 1,222 neurons; each dot = one neuron. **f)** Mean pre-stimulus firing rate for Hit and Miss trials. Asterisk indicates p < 0.05, two-sided paired t-test. Error bars = SEM.

In contrast, neurons that had a greater response to licking than to the whisker stimulus (n = 359), showed no significant difference between Hit and Miss trials when analysed using the same peak-response metric (Supplementary Figure 1; p > 0.05), indicating their activity was not correlated with the behavioural performance. Interestingly, the sensory response profile of these neurons was different from that of neurons whose activity was greatest just after stimulus onset. To compare their sensory response properties independently of lick-related activity, we restricted this analysis to Miss trials, in which no lick response occurred. Neurons with greater stimulus-aligned than lick-aligned activity showed larger sensory-evoked responses and shorter response latencies than neurons with greater peak activity around lick onset (Supplementary Figure 2; p < 0.05; Wilcoxon rank-sum tests). This distinction may help explain why Hit–Miss modulation was evident only among neurons dominated by stimulus-aligned activity, further linking early sensory-evoked SC activity to successful stimulus detection.

### A causal link between superior colliculus activity and behaviour

Our findings so far support the idea that activation of SC neurons contributes to sensory decision-making. To directly test whether this is the case, we optogenetically inhibited SC neurons by activating local GABAergic neurons during the whisker detection task. An AAV containing Cre-dependent ChR2 was injected into the SC in both hemispheres of Vgat-Cre mice, which express Cre recombinase in GABAergic neurons (Supplementary Figure 3a). Mice were then implanted with optic fibres bilaterally in the SC and trained to perform the sensory Go/NoGo task using a whisker stimulus. For these experiments, we used a piezo-driven whisker vibration mesh which allowed us to vary the vibration amplitude (Figure 4a). During the task, blue light (470 nm) was delivered on 50% of randomly selected trials, coinciding with the 1-s whisker vibration. Bilateral photoinhibition of the SC significantly reduced behavioural performance (approximately 50% reduction; Figure 4b-c), with the magnitude of this effect showing a trend with the level of viral expression (Supplementary Figure 3b-c). Detection rates increased with stimulus amplitude, but SC inhibition reduced performance across all amplitudes (Figure 4d; mixed-effects model, p < 0.001). To determine whether inhibiting SC led to slower responses on Hit trials, we analysed reaction times. Mean first-lick reaction time was not significantly altered by SC inhibition (Figure 4e; paired Wilcoxon signed-rank test, p >0.05), with reaction time similar across different stimulus amplitudes (Figure 4f; mixed-effects model, p > 0.05). These findings indicate that inhibition of the SC impaired behavioural performance without slowing the execution of successful responses.

**Figure 4:**
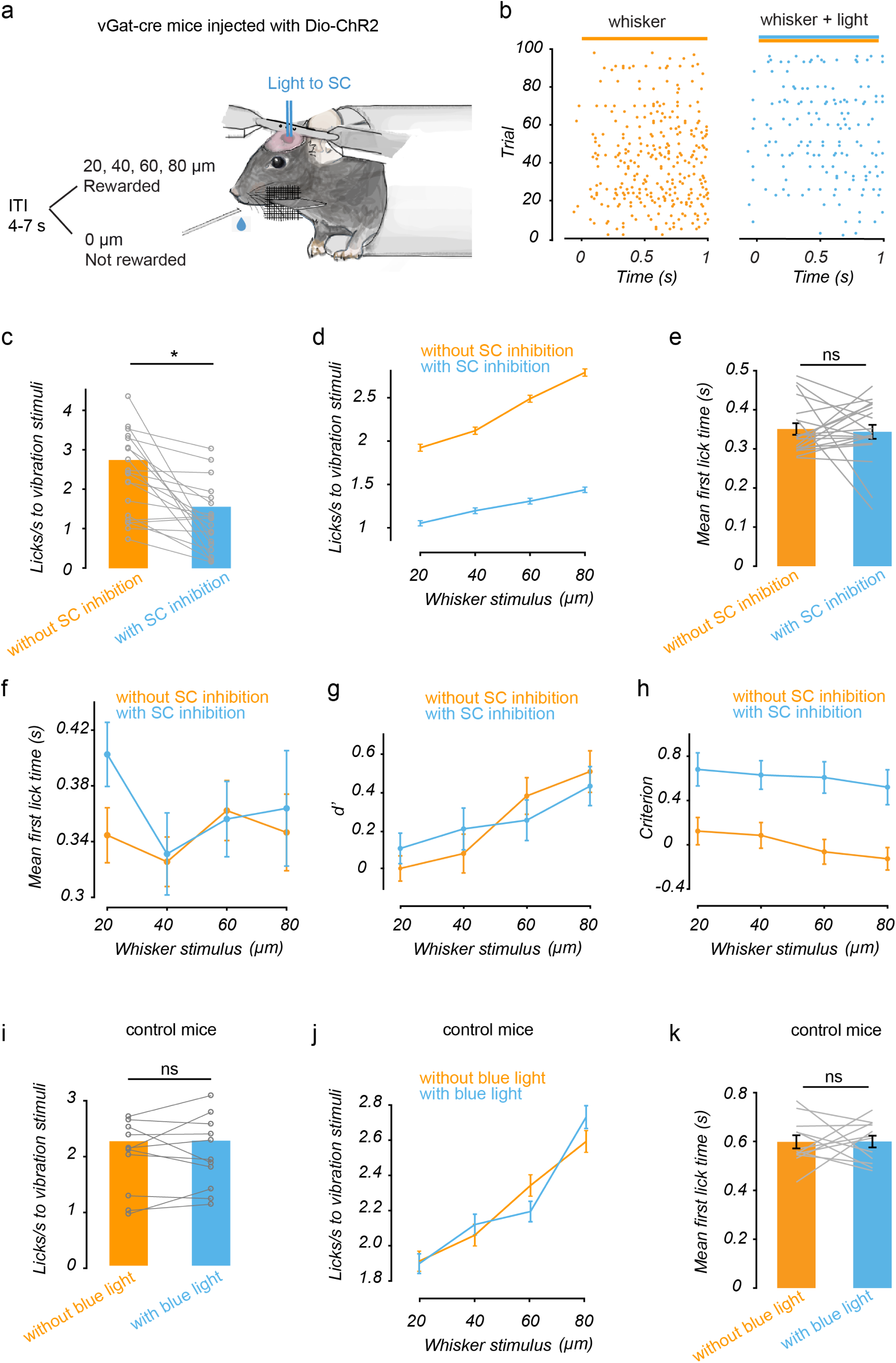
Optogenetic inhibition of the superior colliculus impairs sensory detection performance. **a)** Schematic of the experimental setup. Two optic fibres were implanted bilaterally in the SC of Vgat-Cre mice to activate GABAergic neurons with blue light (470 nm) and hence inhibit the SC. Licking during the 1 s reward window after stimulus onset produced a sucrose reward. Inter-trial interval: 4-7 s. **b)** Example session raster plots with (right) or without (left) optogenetic inhibition. Each dot = a lick; each row = a trial. **c)** Lick rates are plotted without (orange) or with (blue) bilateral photoinhibition (5 mice; 4 sessions each; 20 sessions total). Asterisk indicates p < 0.05, two-sided paired t-test. **d)** Mean lick rate as a function of whisker-vibration amplitude without (orange) and with (blue) SC inhibition (n = 20 sessions). Error bars = SEM. **e)** Reaction times, measured as the latency to the first lick following stimulus onset, are plotted in the presence (blue) or absence (orange) of bilateral photoinhibition. Error bars = SEM. **f)** Mean first-lick reaction time as a function of whisker stimulus amplitude. **g)** Signal detection sensitivity (d′) as a function of whisker stimulus amplitude. **h)** Signal detection criterion as a function of whisker stimulus amplitude. **i)** The effect of light delivery in the control wild-type mice (3 animals; 4 sessions each; 12 sessions total; p > 0.3). **j)** Mean lick rate as a function of whisker-vibration amplitude in control mice without and with blue-light delivery (n = 3 mice; 4 sessions each; 12 sessions total). **k)** Reaction times in control animals (p > 0.9). Error bars = SEM.

To further characterise the effect of SC inhibition on behavioural performance, we quantified signal detection sensitivity (d′) and response criterion ^39^. SC inhibition did not significantly alter d′ across different stimulus amplitudes (Figure 4g; mixed-effects model, p > 0.05), indicating that the reduction in behavioural performance was not accompanied by a detectable change in sensory discriminability. In contrast, SC inhibition increased response criterion (Figure 4h; mixed-effects model, p < 0.001), indicating that animals were less likely to report stimulus presence. Together with the absence of a significant change in reaction time, these findings suggest that SC inhibition primarily altered the likelihood of reporting the stimulus rather than sensory sensitivity or the speed of response execution.

One potential concern with optogenetic manipulations is that scattered light from the optic fibres could activate the retina and indirectly affect performance ^40,41^. To control for this, we implanted optic fibres in wild-type C57BL/6J mice (n=3) and delivered blue light on 50% of trials during whisker stimulation, using the same intensity as in Vgat-Cre animals. Light delivery alone did not affect task performance or reaction times (Figure 4i–k). These control experiments confirm that the behavioural effects observed in Vgat-Cre mice were due to optogenetic inhibition of the SC rather than a non-specific effect of the blue light. Together with our electrophysiological data, these optogenetic experiments demonstrate that neuronal activity in the SC is both correlated with the animal’s decision and required for correct detection during this whisker-dependent sensory decision task.

## Discussion

### SC sensory activity is linked to perceptual decisions

A key challenge in systems neuroscience is to understand how neuronal activity in the brain drives sensory decision-making. To reveal the contribution of the SC to sensory decisions, we directly recorded neuronal activity in the SC as mice performed a binary Go/NoGo sensory detection task. We showed that the activity of SC neurons is correlated with the behavioural outcome: activity was higher when animals correctly detected the whisker stimulus compared to when they missed it. Activity differences between Hit and Miss trials were evident during the early post-stimulus period, supporting a contribution of SC sensory activity to behavioural performance. This interpretation was further supported by our finding that optogenetic inhibition of the SC reduced sensory detection. These results support the view that subcortical circuits within the SC encode task-relevant sensory signals that underlie perceptual decision-making.

Consistent with our findings, recent work using a whisker-based task indicates that SC neurons respond more strongly to high-value (Go) compared to lower-value (NoGo) stimuli ^42^. In that study, reducing the opportunity to make a behavioural response diminished both stimulus selectivity and spontaneous firing rates in the SC, suggesting that task engagement modulates SC activity. In our experiments, we observed a small difference in baseline activity preceding Hit and Miss trials, indicating that internal motivational state may contribute to SC activity and behavioural performance. Neural activity in the somatosensory cortex has also been found to be correlated with perceptual decision-making ^43–46^, suggesting that decision-related activity observed in the SC may be inherited from cortical circuits. One argument against this is that while cortical areas often show distributed representations of sensory, motor, and decision information ^47^, SC neurons exhibit stronger selectivity for behaviourally relevant stimuli than neurons in primary somatosensory cortex ^42^. Furthermore, other work shows that sensory decision making in mice is not disrupted by removal of sensory cortex; mice can learn and perform somatosensory detection tasks despite lesions of the primary somatosensory cortex ^48^ and auditory detection tasks despite lesions of the auditory cortex ^49^. Likewise, mice rely on SC activity rather than cortical activity to detect luminance changes ^50^. Our findings also show that sensory responses in the SC are tightly linked to behavioural outcome. Consistent with this, a study in non-human primates performing visual decision-making tasks showed that SC activity reflects factors influencing perceptual decisions and that manipulating SC activity can influence the capacity to report the presence or absence of a stimulus ^51^. Together, these findings suggest that SC neurons are likely to integrate sensory and motivational signals to bias action selection.

### SC activity is required for sensory decision-making

To determine whether SC activity is causally required for performance, we transiently inhibited SC neurons optogenetically during the behavioural task. Inhibition significantly reduced detection accuracy, demonstrating that neuronal activity in the SC directly contributes to somatosensory perceptual decision-making. Importantly, this impairment was accompanied by an increase in response criterion, while perceptual sensitivity (d′) and reaction times showed no detectable change. This pattern suggests that SC inhibition primarily altered the likelihood of reporting stimulus presence, rather than reducing sensory discriminability or slowing the execution of successful responses. This result parallels findings during visual decision tasks, in which transient SC inhibition specifically during the stimulus period, but not during the preparatory or motor phases, impairs detection ^52,53^. Similarly, bilateral inactivation of the SC disrupts performance on cognitively demanding orienting tasks, further supporting its role in selective attention and executive control ^54,55^. Collectively, these observations show that SC activity is both correlated with and necessary for sensory decisions, and that its influence is temporally aligned with sensory processing rather than with downstream motor execution.

In non-human primates, the SC has also been shown to play a causal role in decision-making during a visual perceptual task and during covert attention ^56–58^. Other work suggests that the SC represents the behavioural priority of visual stimuli, influencing perceptual judgements even without overt motor actions ^21,26,57^. Unilateral SC inactivation biases perceptual judgements toward the contralateral field ^56,59^, whereas micro-stimulation enhances attention without evoking eye movements ^60^. These findings show that detection and choice behaviour in non-human primates also depend on activity in the SC during perceptual decisions. Together, this work supports a broader role for the SC in linking sensory information to behavioural choice, rather than simply playing a role of movement generation.

A closely related role for the SC in setting decision criterion has been demonstrated in non-human primates. Crapse and colleagues showed that SC activity tracked changes in decision criterion independently of perceptual sensitivity, and that manipulation of SC activity causally shifted criterion ^51^. Using visual stimuli and saccadic reports, SC stimulation also shifted criterion without a detectable change in reaction time, supporting the idea that the SC can influence whether a stimulus is reported without necessarily altering response latency. Here, we observed a comparable dissociation between criterion and sensitivity during somatosensory detection reported by licking. This raises the possibility that SC contributions to decision criterion may generalise across sensory modalities, species and types of behavioural response. Decision-related changes in response criterion have also been linked to activity in premotor cortex during vibrotactile detection in non-human primates ^61^, suggesting that criterion setting may involve interactions between cortical and subcortical decision circuits. In contrast, a visual detection study in mice found that unilateral SC inhibition increased reaction times for contralateral visual stimuli ^53^, suggesting that the effects of SC inhibition on response timing may depend on the specific task and experimental context. Many models of perceptual decision-making propose that sensory evidence is accumulated over time until sufficient evidence is available to make a choice ^62,63^, and such models have also been applied specifically to Go/NoGo decisions ^64^. Changes that slow evidence accumulation in these models would be expected to be reflected in longer decision times. The preservation of d′, lack of a detectable change in reaction time, and increase in criterion suggest that SC inhibition predominantly altered the animals’ likelihood of reporting stimulus presence, without a detectable reduction in sensory discriminability or slowing of successful responses. This is consistent with a role for the SC in regulating how sensory evidence is translated into behavioural choice.

### Circuit mechanisms supporting SC-dependent decisions

Our findings raise the question of how SC circuits impact sensory decision-making. Recent work shows that neuronal activity in the SC reflects the convergence of cortical and subcortical pathways within sensorimotor zones ^31^. Whisker-related regions within the mouse SC form a hub that integrates somatosensory and motor cortical signals via parallel excitatory and inhibitory pathways ^31,65^. The SC also participates in reciprocal interactions with frontal cortical circuits, through which SC activity can modulate competing behavioural choice signals during decision-making ^23^. In somatosensory-guided action selection tasks, cooperative interactions between GABAergic and glutamatergic neurons in the SC drive choice-related activity in frontal cortical areas via thalamic pathways, supporting reciprocal communication between sensory and executive systems ^23^. Together, these SC circuits are likely to support the coordinated sensory–motor transformations required for sensory decision-making.

Beyond these circuit interactions within the somatosensory system, the SC also contributes to broader decision-related functions. In rodents, SC neurons encode both task context and spatial choice during visually guided decisions, with choice-related signals emerging earlier in the SC than in frontal cortical regions such as the prefrontal cortex ^54^. Consistent with this, neural activity in the mouse SC is enhanced at cued spatial locations rather than suppressed at uncued ones, indicating a bias toward behaviourally relevant stimuli ^66^. Recent work further shows that sensory and attention-related responses in the mouse SC are strongly shaped by learned behavioural relevance, with task-related modulation particularly prominent in the deeper SC layers ^67^. Together, these findings suggest that SC representations are shaped not only by sensory input itself, but also by the behavioural significance assigned to that input through task demands and experience.

In addition to cortical and thalamic interactions, the SC also forms reciprocal connections with basal ganglia structures that regulate action selection. The substantia nigra pars reticulata provides a major inhibitory input to the SC, controlling orienting and target selection ^68–70^. Other work shows that activation of the direct striatal pathway modulates SC neuronal activity and alters performance in sensory detection tasks ^71^. In parallel, the basal ganglia can both enhance and suppress sensory activity in the SC through complementary mechanisms that are thought to support the selection of behaviourally relevant targets while suppressing distractors ^72^. SC neurons projecting to dopamine and GABA neurons in the ventral tegmental area and substantia nigra pars compacta also contribute to cue-guided learning through dopaminergic modulation ^73^. Notably, inhibitory basal ganglia input to the SC has been proposed as a potential mechanism for setting decision criterion by providing a bias signal that influences whether sensory evidence leads to a behavioural response ^51^. Related computational work has proposed a cortico–basal ganglia–SC circuit in which the SC detects threshold crossing, while the cortico–basal ganglia pathway adaptively tunes the decision threshold ^74^. Together, these proposals provide a potential circuit framework through which basal ganglia–SC interactions could influence whether accumulated sensory evidence is translated into a behavioural response. More broadly, the SC-basal ganglia interactions may provide a mechanism for integrating sensory evidence with reinforcement signals to guide adaptive behaviour ^75,76^.

In conclusion, our results demonstrate that the mouse SC plays a causal role in transforming whisker-derived sensory signals into behavioural decisions. This extends the traditional view of the SC as a reflexive orienting centre and positions it as a key node within a distributed network supporting perceptual decision-making. The selective increase in response criterion, in the absence of detectable changes in perceptual sensitivity or reaction time, suggests that the SC influences whether sensory information is translated into a behavioural report, rather than simply determining sensory discriminability or response speed. Our findings therefore extend evidence for criterion-related functions of the SC from visual decisions reported by eye movements in primates to somatosensory decisions reported by licking in mice. Future work combining population-level recordings with pathway-specific perturbations will be essential to determine how SC representations dynamically interact with cortical decision networks over time.

## Methods

### Animals

A total of 7 adult male C57BL6/J and 5 adult male Vgat-ires-Cre mice (The Jackson Laboratory) were used in this study. Mice were 4-7 weeks old at the start of the experiment. Animals were housed in a controlled environment with a 12-hour reversed light-dark cycle at a temperature of 22°C. All animal procedures were approved by the Animal Experimentation Ethics Committee of the Australian National University. Mice were water-restricted, and their body weight and health were monitored daily. Animals received free access to water for 2 hours after each training session, and food was available *ad libitum*.

### Surgery for the recording experiments

C57BL6/J animals were placed in a chamber to induce light anaesthesia via brief exposure to isoflurane (3.5% in oxygen), then mounted in a stereotaxic frame with anaesthesia continued using isoflurane (1-1.5% in oxygen) delivered through a nose cone. Throughout surgery, mice were placed on a servo-controlled heating blanket (Harvard Instruments) to maintain a steady body temperature near 37°C. Atropine (0.3 mg per kg, 10% weight per volume in saline) was administered subcutaneously to reduce secretions. The eyes were covered with a thin layer of Viscotears liquid gel (Alcon, UK). A craniotomy (diameter 3 mm) was performed above the left SC (0.5 mm anterior to lambda and 1.5 mm lateral to midline) while keeping the dura intact. The craniotomy was covered using a 3 mm glass coverslip (0.1 mm thickness, Warner Instruments, CT). The coverslip was glued to the bone surrounding the craniotomy. The mice were then implanted with a custom-designed head-bar using dental cement anterior to the cranial window. A thin layer of a silicon sealant (Kwik-Cast, World Precision Instruments, USA) was applied to cover all parts of the cranial window and the skull. Following the surgery, ketoprofen (5 mg per kg) was given for pain relief.

### Behavioural task for electrophysiological recordings

Mice implanted with a head-bar were allowed to recover from surgery for 1 week and then placed on a water restriction schedule, receiving unrestricted water for 2 hours daily. They were then gradually habituated to the experimenter to the head-fixation setup. During head fixation, mice were placed on a circular disc, which they could turn freely by running (Figure 1). The eyes and the whiskers were also constantly monitored via two high-speed video cameras. When the mice were habituated to the head fixation, the first stage of training began, during which the animals received a 5% sucrose reward for every lick from a custom 3D-printed “lick-port” connected to an Arduino UNO board (Duinotech Classic, Cat#XC4410). Once animals reliably licked to receive a sucrose reward, they were moved to the next stage of training, in which they were rewarded only when presented with a sensory stimulus (“Go” trials). Whisker stimulation was delivered via an air-puff to the right vibrissal pad (contralateral to the craniotomy for recording). The air-puff stimulus was administered through a rubber tube connected to a 12V gravity-fed solenoid valve, which controlled airflow from a pressurised air supply (∼10 psi) through the valve opening. The rubber tube was aimed at an angle of approximately 70° relative to the horizontal to target the middle region of the mouse’s whiskers. The distance of the air-puff tube from the mouse and its tip diameter were adjusted so that air puffs fully engaged all whiskers on the right side of the snout. White noise (∼60 dB SPL) was used to mask the sound produced by solenoid activation. During catch trials, there was no air-puff. Licking during catch trials resulted in no reward (“NoGo” trials). In the final stage of the task, the air-puff stimulus had a duration of 500 ms and an inter-trial interval of 7-13 s (∼200 trials per session). “Hit” trials were defined as trials where there was at least one lick 0.05-1.5 s post stimulus onset. “Miss” trials were defined as the absence of licking 0.05-1.5 s post stimulus onset.

Behavioural performance was quantified using the Hit rate and False Alarm rate. Behavioural sensitivity (d′) was calculated as the difference between the *z*-transformed Hit and False Alarm rates. To avoid infinite values, Hit and False Alarm rates were corrected using the log-linear correction before calculating d′. Behavioural discrimination was also quantified using the area under the receiver operating characteristic curve (AUROC), calculated from binary behavioural responses during stimulus and catch trials. Mean d′ and AUROC values were compared with chance levels (0 for d′ and 0.5 for AUROC) across recording sessions using two-sided Wilcoxon signed-rank tests.

### Neuropixel recording and analysis

We recorded neuronal activity in the SC as mice performed the detection task described above. Extracellular single-unit activity was recorded with high-density Neuropixel 1.0 probes, allowing hundreds of neurons to be recorded simultaneously across multiple layers of the anterolateral region of the SC. Each 1 cm long Neuropixel probe has 960 electrode sites with 384 active channels. Signals from all 384 channels were simultaneously amplified, filtered (250-5,000 Hz) and continuously recorded onto disk at a sampling rate of 30 kHz. Before the first Neuropixel probe insertion, the glass coverslip covering the craniotomy was removed. The probe was then inserted into the SC and allowed to settle for ∼10–20 min before the start of the recordings. Between sessions, the probe was withdrawn, and the craniotomy was sealed with sterile silicone (Kwik-Cast) and removed before the next recording session. Data were sorted off-line using the Kilosort package (https://github.com/MouseLand/Kilosort), followed by manual curation with the Phy GUI (https://github.com/cortex-lab/phy). To categorise spike clusters as single units, we assessed their spike waveforms, cross-correlograms and electrode positions, making adjustments by merging or splitting clusters when needed.

For comparison of firing rates during Hit and Miss trials, we used a short 100-ms analysis window after stimulus onset. For these comparisons, only sessions that contained at least 35 Hits with at least one lick 0.05–1.5 s post stimulus onset per trial were selected. Response latency was defined as the first occurrence, after stimulus onset, of two consecutive bins in the peri-stimulus time histogram (PSTH; 5 ms bin width) where there was a significant increase in spike count (p < 0.05; t-test).

For the analyses presented in Figure 2, stimulus-aligned responses were calculated from Hit trials, whereas lick-aligned responses were calculated relative to the first rewarded lick occurring 0.05–1.5 s after stimulus onset. Neurons were grouped according to whether their maximum firing rate within ±100 ms of stimulus onset was greater or lower than their maximum firing rate within ±100 ms of first-lick onset. For the spontaneous-lick control analysis, neuronal activity was aligned to spontaneous licks occurring during the pre-stimulus period (−3 to 0 s) in Hit trials.

To compare sensory response properties between neurons with greater stimulus-aligned than lick-aligned activity and neurons with greater lick-aligned than stimulus-aligned activity, analyses were restricted to Miss trials to avoid contamination from lick-related activity. Firing rates were baseline-subtracted using the 100-ms period preceding stimulus onset, and mean and peak sensory responses were quantified within the first 100 ms following stimulus onset. Response latency was calculated as described above. Sensory response properties were compared between the two neuronal populations using two-sided Wilcoxon rank-sum tests.

Statistical significance was set at p < 0.05. Results are presented as average values ± the standard error of the mean (SEM). In the figures, “ns” denotes not statistically significant, whereas an asterisk denotes p < 0.05.

### Surgery for the optogenetic experiments

Glass pipettes (Drummond), pulled on a microelectrode puller (Sutter Instrument Co.; P-87, USA) and broken to give a diameter of around 20 μm, were back-filled with mineral oil and front-loaded with viral suspension (AAV1-Ef1a-DIO-hChR2(E123A)-EYFP; University of Pennsylvania, USA). Vgat-ires-Cre or C57BL/6J control mice were anaesthetised with isoflurane (induction 3.5%, maintenance 1–1.5%) and maintained at 37 °C. Two craniotomies with a diameter of 1 mm were performed above both sides of the SC (0.5 mm anterior to lambda and 1.5 mm lateral to midline). AAV1-Ef1a-DIO-hChR2(E123A)-EYFP was then injected into intermediate/deep layers of SC of the Vgat-cre mice (1.5-2.0 mm from the surface; 100-180 nl; 36.8 nl per minute; Nanoject II, Drummond). After injections, we implanted optic fibres (125 µm core diameter; Thorlabs) bilaterally using dental cement in both Vgat-cre and control mice. The optic fibre was inserted vertically into the SC (1.5 mm from the surface). The mice were then implanted with a custom-designed head plate using dental cement anterior to the implanted optic fibres. Ketoprofen (5 mg/kg; subcutaneous) was given for pain relief. After surgery, mice were returned to their cage and placed on a heating pad to recover.

To confirm viral expression and optic fibre location, at the end of behavioural experiments, animals were perfused transcardially with 0.9% sodium chloride solution and then 4% paraformaldehyde (PFA). The brain was removed from the skull and kept in PFA overnight. Coronal slices (50 μm thick) were prepared, DAPI-stained and examined under a confocal microscope (Zeiss LSM800 with Airyscan). For the quantification of fluorescence between different mice, the same microscope settings were used across different animals.

### Optogenetic behavioural experiments and analyses

Following recovery, mice were water-restricted and habituated to an acrylic head-fixation tube. A capacitive lick-port was positioned ∼0.5 mm below the lower lip and ∼5 mm posterior to the snout. Whisker deflections were applied to the right vibrissal pad using a light-weight fine mesh plate glued to a piezoelectric ceramic (Morgan Matroc, Bedford, OH). The piezo was driven by an amplifier (PiezoDrive, amplification gain of 20) controlled by commands generated in MATLAB and sent to the analogue output of a PCIe-6321 data acquisition board (National Instruments; 20 kHz sampling rate). Five vibration amplitudes (0, 20, 40, 60 and 80 µm) were delivered in the vertical direction with a variable inter-trial interval of 4–7 s. The vibration duration was 1 s. Vibration amplitudes were pseudo-randomised in blocks of 5 trials, with each block having all amplitudes (∼200 trials per session).

After training, the implanted optic fibres were connected to a 470 nm LED (Thorlabs), with the power of blue light out of the fibre tip being ∼1.8 mW. The LED was controlled through a National Instruments board using scripts written in MATLAB. The whisker vibration stimuli were presented either with or without optogenetic inhibition of the SC (10 stimulus conditions in total; the optogenetic inhibition in 50% of the trials). In all experiments, whisker stimulation was presented simultaneously with optogenetic activation of the SC (duration, 1 s).

Behavioural analysis included the first four sessions after introducing inhibition trials. Reaction time was calculated only for Hit trials and was defined as the latency from stimulus onset to the first lick occurring within the response window. To assess whether SC inhibition altered perceptual sensitivity or response strategy, signal detection measures were calculated for each mouse-session and stimulus amplitude (20, 40, 60 and 80 μm). Hit rate was calculated as the proportion of Go trials with a lick response, and false-alarm rate as the proportion of lick responses during catch (0 μm) trials. Hit and false-alarm rates were adjusted using the log-linear correction to avoid infinite z-scores. Signal detection sensitivity (d′) was calculated as *d*’ = *Z*(Hit rate) − *Z*(False alarm rate), and response criterion was calculated as *c* = −0.5 × [*Z*(Hit rate) + *Z*(False alarm rate)], where *Z* denotes the inverse cumulative normal distribution ^39^.

Data were analysed using MATLAB scripts. Pairwise comparisons were performed using two-sided paired t-tests and confirmed using Wilcoxon signed-rank tests. Behavioural measures that varied across stimulus amplitude (lick rate, reaction time, d′ and response criterion) were analysed using linear mixed-effects models, with optogenetic inhibition and stimulus amplitude as fixed effects and random intercepts for mouse and mouse-session to account for repeated measurements.

## Acknowledgments

This work was supported by the Australian Research Council (ARC) Discovery Project Grant (DP240103043), the ARC Centre of Excellence for Integrative Brain Function (CE140100007) awarded to G.J.S and E. A., and the Australian National University and Monash University.

## Author Contributions

S.G., E.A. and G.J.S. designed the experiments and interpreted the data. S.G. and M.T. performed and analysed the Neuropixel experiments. S.G. performed and analysed the optogenetics experiments. S.G. drafted the manuscript, and all authors edited and approved the final version of the manuscript.

## Competing Interests Statement

The authors declare no competing interests.

## Data and code availability

The data that support the findings of this study are available from the corresponding authors upon reasonable request. This paper does not report original code.

## Supplementary Figures

**Supplementary Figure 1:**
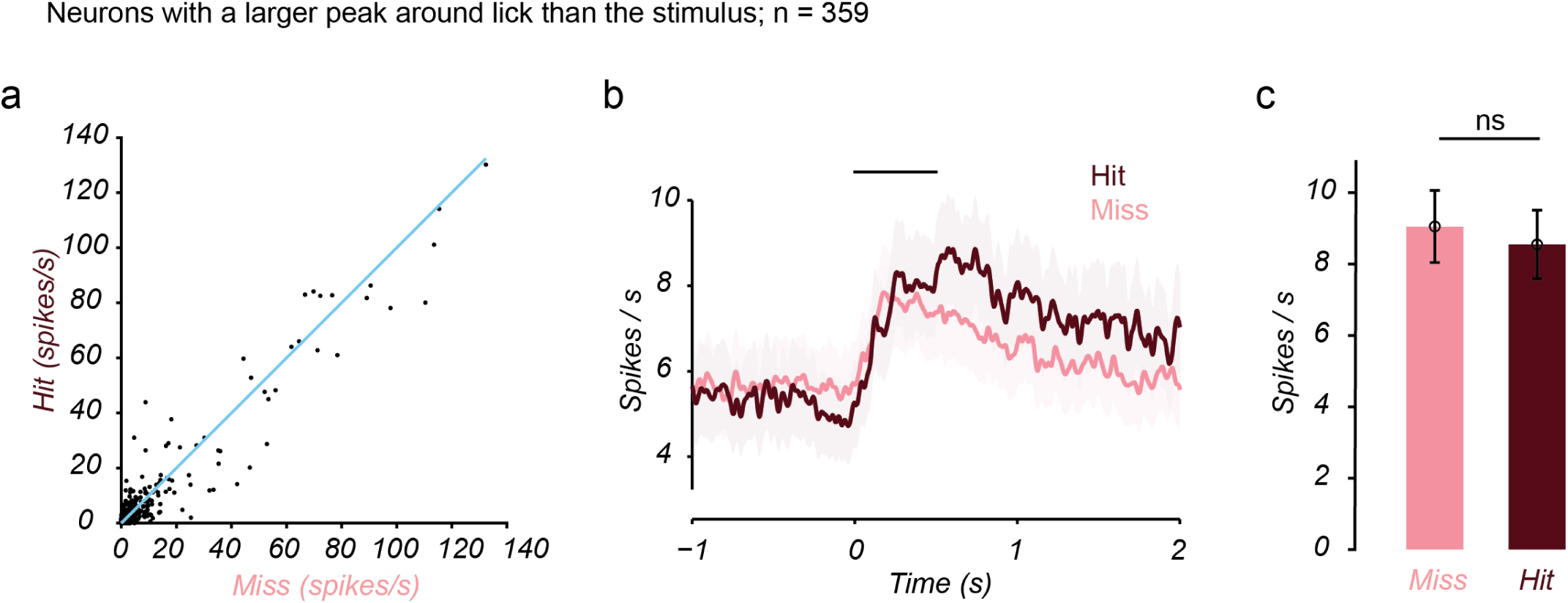
Hit and Miss responses in neurons with greater lick-aligned than stimulus-aligned activity. **a)** For each neuron, we plotted the maximum firing rate within the first 100 ms following stimulus onset for Hit and Miss trials. For these analyses, only neurons whose activity was greater around lick onset than around stimulus onset were included (359 neurons). **b)** Mean population activity over time for Hit (burgundy) and Miss (pink) trials. Time 0 = stimulus onset; black line = 0.5 s stimulus period. As in panel **a,** only neurons with greater lick-aligned than stimulus-aligned activity are plotted. **c)** Bar graph showing the average peak firing rate of these neurons within the first 100 ms following stimulus onset for Hit and Miss trials. Error bars = SEM.

**Supplementary Figure 2:**
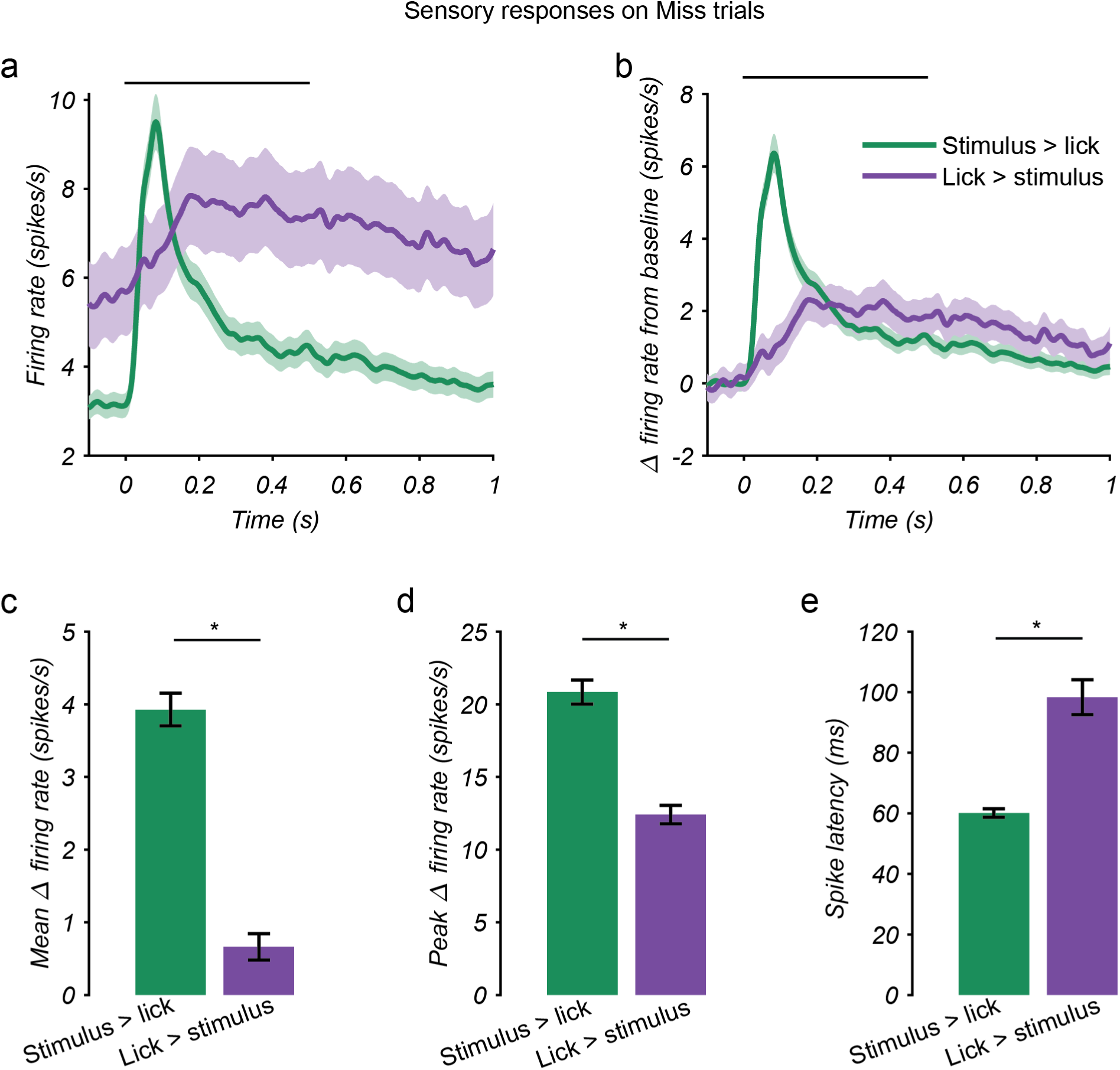
Sensory response profiles on Miss trials for neurons grouped by the relative magnitude of stimulus- and lick-aligned activity. **a)** Mean firing rate on Miss trials aligned to whisker stimulus onset for neurons with greater stimulus-aligned than lick-aligned activity (green; n = 1,222) and neurons with greater lick-aligned than stimulus-aligned activity (purple; n = 359). The black line indicates the stimulus period. Shaded areas represent ± SEM. **b)** Baseline-subtracted mean firing rate on Miss trials for the same neuronal populations as in panel **a**. Shaded areas represent ± SEM. **c)** Mean baseline-subtracted firing rate during the first 100 ms following stimulus onset. **d)** Peak baseline-subtracted firing rate during the first 100 ms following stimulus onset. **e)** Response latency following stimulus onset for the same neuronal populations. Between-group comparisons were performed using two-sided Wilcoxon rank-sum tests. Asterisks indicate p < 0.05. Error bars = SEM.

**Supplementary Figure 3:**
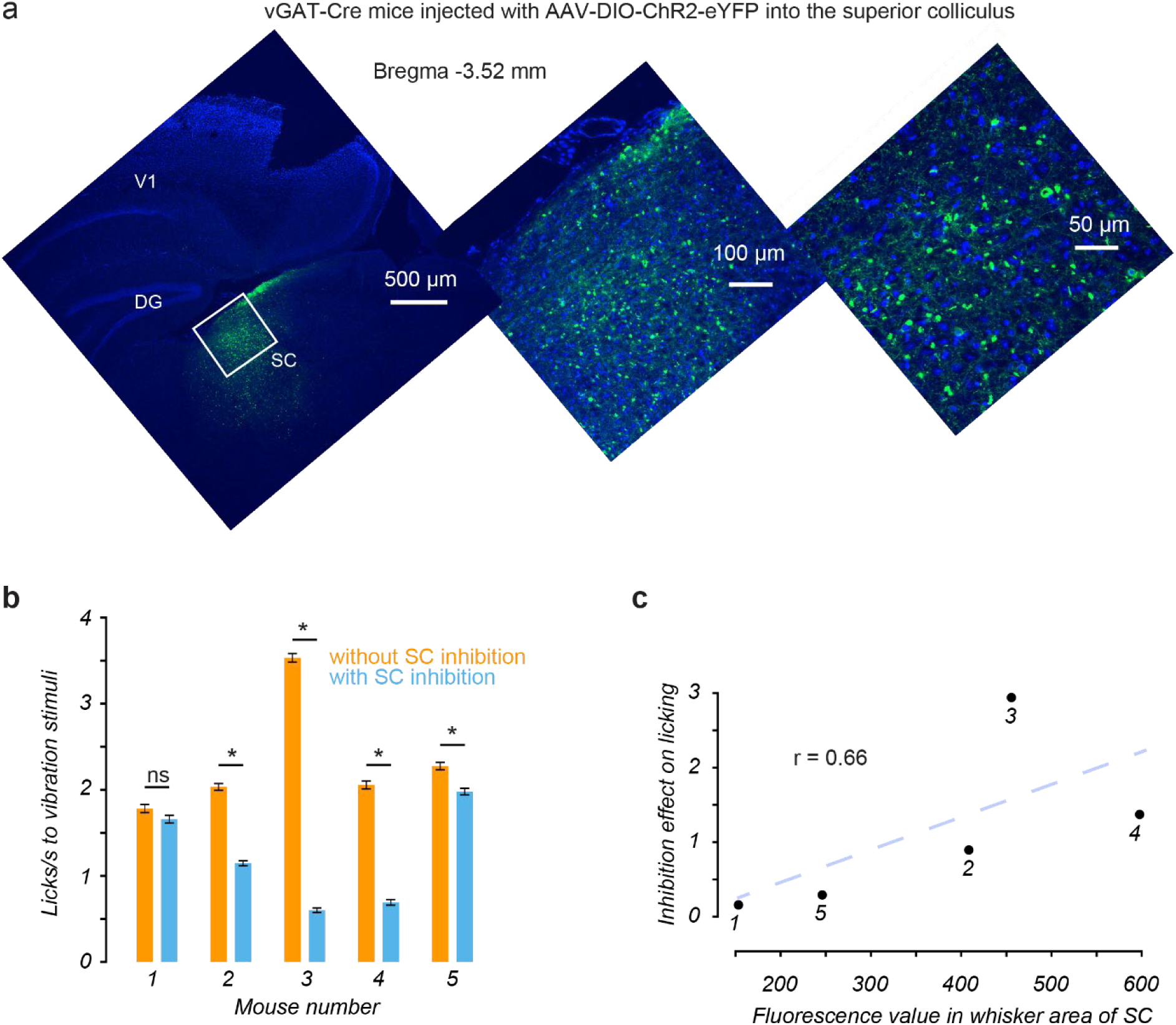
Viral expression and its relationship to behavioural effects. **a)** Coronal section showing the viral injection site in the SC. eYFP fluorescence is shown in green, and DAPI in blue. **b)** Bilateral photoinhibition of the SC significantly reduced the lick rate in 4 out of 5 mice (4 sessions each). Asterisks indicate p < 0.05 (two-sided paired t-test). **c)** The magnitude of photoinhibition for each mouse is plotted against the measured level of viral expression.

## Notes

### Competing Interest Statement

The authors have declared no competing interest.

### Summary of Updates

The manuscript has been revised to include additional analyses of behavioural performance, including signal detection sensitivity, response criterion, and reaction time, and additional analyses comparing sensory response properties of neuronal populations. Figure 4 has been updated and a new Supplementary Figure 2 has been added. The Results, Methods, and Discussion have been updated accordingly.

